# Immune pathways that regulate neutrophil activation and replenishment prevent persistent pneumococcal colonization

**DOI:** 10.64898/2026.01.15.699644

**Authors:** Teniola Idowu, Kristen Lokken-Toyli, Stephen Yeung, Jeffrey N Weiser

## Abstract

Here, we examine immune signaling pathways affecting the duration of primary nasopharyngeal colonization by *Streptococcus pneumoniae* (*Spn*), the first step in its pathogenesis. *Spn* colonization which lasts days to weeks in WT mice was persistent (≥ 6 months) in the absence of IL-17RA-signaling. RNA-seq analysis confirmed the role of IL-17RA signaling in neutrophil-associated pathways. Rather than driving neutrophil specific chemokine expression, IL-17RA-signaling was required to replenish neutrophils in nasal tissue that were otherwise depleted during infection. Enhanced neutrophil trafficking correlated with IL-17RA-dependent expression of endothelial cell adhesion molecules that promote neutrophil trafficking from the circulation into nasal tissue. Persistent colonization was also observed in mice lacking IL-1R-signaling. Recognition of IL-1-family cytokines, however, was not necessary for the expression of IL-17A or neutrophil recruitment. Instead, IL-1R-signaling was associated with the activation of neutrophils in nasal tissue that displayed increased levels of the surface marker CD11b, an important receptor for the complement-opsonized phagocytosis of *Spn*. Our findings provide insight into the requirement for sustained neutrophil presence and activity to prevent persistent mucosal infection by a leading opportunistic mucosal pathogen.

**Author Summary:** Colonization of the upper airway by Streptococcus pneumoniae (*Spn*) is common and often asymptomatic, particularly in early life, yet prolonged colonization increases the risk of transmission and invasive disease. Despite its importance, the immune mechanisms that prevent long-term persistence of *Spn* in the upper airway are not fully understood. In this study, we investigated how two immune signaling pathways, the IL-17RA and IL-1R pathways, contribute to preventing persistent *Spn* colonization. Using a mouse model, we found that the IL-17RA pathway supports continued replenishment of neutrophils into nasal tissue by promoting expression of molecules that allow these cells to exit the bloodstream and enter the nasal tissue. In contrast, the IL-1R pathway activates neutrophils in a way that makes them more effective at clearing *Spn*. Loss of either pathway resulted in persistent *Spn* colonization, with an additive effect seen when both pathways were removed. Collectively, our results highlight how two distinct immune pathways cooperate to control pneumococcal colonization by sustaining neutrophil recruitment and function.

## Introduction

Many members of the extensive and diverse microbial flora appear to stably colonize mucosal surfaces. For other microbes, including leading opportunistic pathogens, colonization is transient. The duration of colonization plays a key role in the epidemiology of many leading infectious diseases. Longer episodes of carriage extend the risk of more invasive infections from the colonizing reservoir and provide a greater opportunity for transmission to new hosts.

We have studied the dynamics of colonization using *Streptococcus pneumoniae* (*Spn*), a leading opportunistic bacterial pathogen of the human respiratory tract. *Spn* resides on the mucosal surfaces of the human nasopharynx [1]. Natural carriage of *Spn* is characterized by sequential episodes of colonization with each lasting from days to months [2]. Both the prevalence and duration of colonization tend to be higher during infancy – an observation that correlates with the increased incidence of invasive *Spn* disease during early life.

IL-17RA signaling, in particular, has been shown to play a key role in controlling natural *Spn* carriage and disease. For example, gene polymorphisms associated with IL-17A levels are correlated with the *Spn* colonization rate in young children [3]. In population studies of children, a higher density of colonizing *Spn* is associated with lower IL-17A production by PBMCs [4]. IL-17A can be elicited by *Spn* stimulation of tonsillar cells from children [5]. Experimental human carriage in healthy adults leads to increased numbers of pulmonary IL-17^+^CD4^+^ T-cells [6]. The effects of IL-17RA signaling have also been extensively studied in murine models of colonization and disease. In mice, clearance of primary colonization does not require *Spn*-specific antibody and depends on CD4^+^ T cell driven immunity [7]. Vaccine-mediated induction of IL-17RA cytokines by Th17 cells accelerates clearance within days, and inhibition of the IL-17 responses decreases colonization density [5],[8]. Additionally, adoptive transfer of CD4^+^ T cells from *Spn* immune mice protects against infection in an IL-17A-dependent manner [9]. On the other hand, no association of colonization dynamics and Th17 cells was found in a study of experimental human pneumococcal carriage, although the limited duration of 10 days in this model might be too brief to observe any effect [10].

What is less clear is how Th17 immunity and the production of IL-17 family cytokines impact colonization [11],[12]. In general, IL-17 protective responses are thought to be mediated through the increased expression of chemokines that enhance the recruitment of neutrophils to the site of infection [13]. *Spn* colonization, however, is characterized by a mild acute inflammatory response in which neutrophils in the nasal lumen are observed only during the first week of infection rather than during later weeks when clearance is most robust [8]. An important role for neutrophils in *Spn* clearance during colonization is highly plausible considering that, as an encapsulated pathogen, effective host defense requires uptake and killing by professional phagocytes following bacterial opsonization. It has also been suggested that IL-17 family cytokines may have a direct effect on phagocytic cells by promoting the uptake and killing of opsonized *Spn* [5],[6]. Another potential factor in the interaction of *Spn* and neutrophils is the expression of the pore-forming toxin, pneumolysin, capable of lysing host cells following bacterial engulfment [14],[15].

In addition to IL-17RA signaling, clearance of colonization involves recognition of IL-1R family cytokines [16]. The IL1R is important in initiating clearance in adult mice, although it has no effect on early colonization of infant mice, which show reduced baseline expression of genes involved in IL-1R signaling [17]. IL-1-signaling appears to depend on the pneumolysin-mediated release of IL-1α, and intranasal treatment with IL-1α is sufficient to reduce *Spn* colonization density. It is unclear if and how these two pathways, involving IL-17RA and IL-1-signaling, coordinate to facilitate the eventual clearance of colonization.

The overall goal of this study is to leverage the mouse model to define how immune signaling pathways affect mucosal immunity to limit the duration of the prolonged colonization that is typical of *Spn*.

## Results

### IL-17RA-dependent signaling prevents persistent colonization

To investigate the role of IL-17RA signaling in *Spn* clearance, we compared colonization dynamics in wild-type (WT) and IL-17RA receptor knockout (*Il17ra^-/-^*) mice. In infant mice **(Fig 1A)**, WT animals gradually reduced bacterial density and fully cleared colonization by 9 weeks post-infection (p.i.). In contrast, *Il17ra^-/-^* mice remained colonized throughout the study, with high colonization levels persisting through 26 weeks p.i., at which point we considered them as persistently colonized. During this period, colonized mice grew and behaved normally. Infants are known to have prolonged carriage duration, so we determined if we see this same effect in adult mice [18]. To assess this, the experiment was repeated in adult mice 6-8 weeks of age **(Fig 1B)**. WT adults cleared colonization within 3 weeks, while *Il17ra*^-/-^ adults, similar to *Il17ra^-/-^*infants, remained persistently colonized. These findings demonstrate that IL-17RA-signaling is essential for promoting clearance and preventing persistent colonization- an effect that is independent of host age.

**Fig 1:**
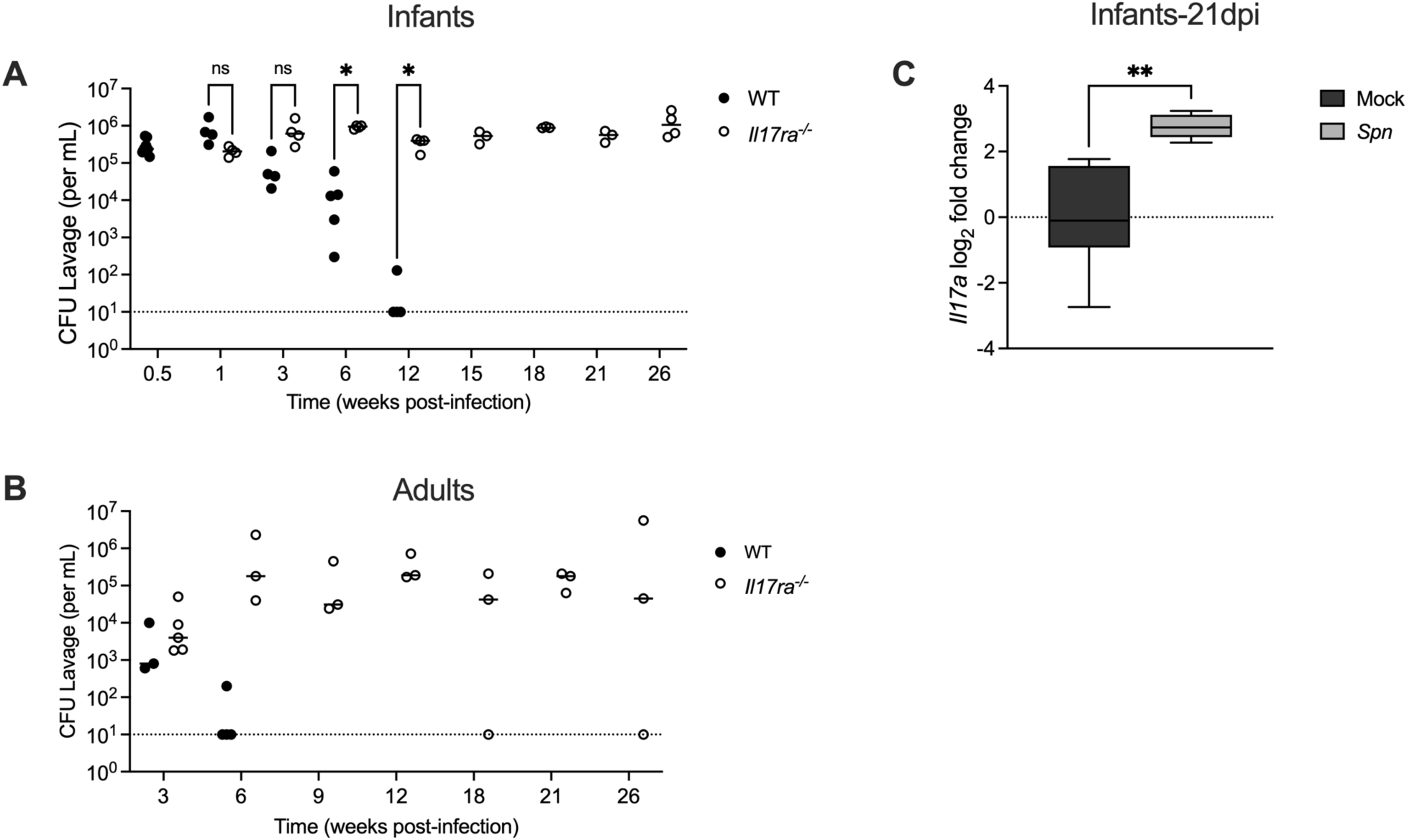
IL-17RA signaling prevents persistent colonization. Wild type (WT) and IL-17RA-deficient (*Il17ra^-/-^*) mice were colonized intranasally with serotype 23F *S.pneumoniae* (*Spn*). Nasal lavages were collected to quantify bacterial colonization density (CFU/mL). Colonization density was assessed at the indicated timepoints in (**A)** infant mice colonized with *Spn* at 4-7 days old (n=4-8) and (**B**) adult mice colonized with *Spn* at 6-10 weeks of age (n=3-5). Dotted line represents the lower limit of detection. Each circle represents an individual mouse and the horizontal line denotes the median for each group. Statistical significance determined using multiple unpaired Students *t* tests. Significance is designated by * = p< 0.05; ns = not significant. **(C)** At 21 days post-infection (dpi) mock- and *Spn*-colonized WT infant mice were euthanized and RNA isolated from nasal lavages was used to quantify *Il17a* expression by qRT-PCR. Expression was normalized to *Gapdh* and calculated as fold change relative to mock-infected controls. Each group represents 5-6 mice and data are presented as a box-and-whisker plot showing the median, interquartile range, and min–max values. Statistical significance determined using an unpaired Students *t* test. Significance is designated by ** = p < 0.01.

Next, to confirm activation of the IL-17 pathway during clearance, we measured expression of the cytokine IL17A by qRT-PCR in nasal tissue of mock and colonized WT infant mice **(Fig 1C)**. Colonized WT infants exhibited an increase in *Il17a* expression at 21-days post-infection (dpi), which coincides with the onset of *Spn* clearance.

### Colonization induces robust immune responses in wild-type but not *Il17ra^-/-^* mice

To determine what host pathways IL-17RA-signaling affects to prevent persistent colonization, we performed RNA-sequencing on transcripts isolated from nasal lavage samples followed by Reactome pathway enrichment analysis [19]. In WT mice, colonization triggered robust upregulation of immune-related pathways, including those involved in innate defense and neutrophil function **(Fig 2A)**. In contrast, *Il17ra^-/-^*mice exhibited a blunted transcriptional response with many pathways having minimal activation with few key immune processes reaching statistical significance **(Fig 2A)**. Notably, the neutrophil degranulation pathway was among the most highly enriched pathways in WT mice, but was not significantly upregulated in *Il17ra^-/-^* mice. Multiple individual genes in this pathway showed increased expression in response to colonization in WT mice, but not in *Il17ra^-/-^* mice **(Fig 2B)**. These findings suggest that IL-17RA signaling is required for full activation of host immune programs during colonization, especially pathways critical for neutrophil responses.

**Fig 2:**
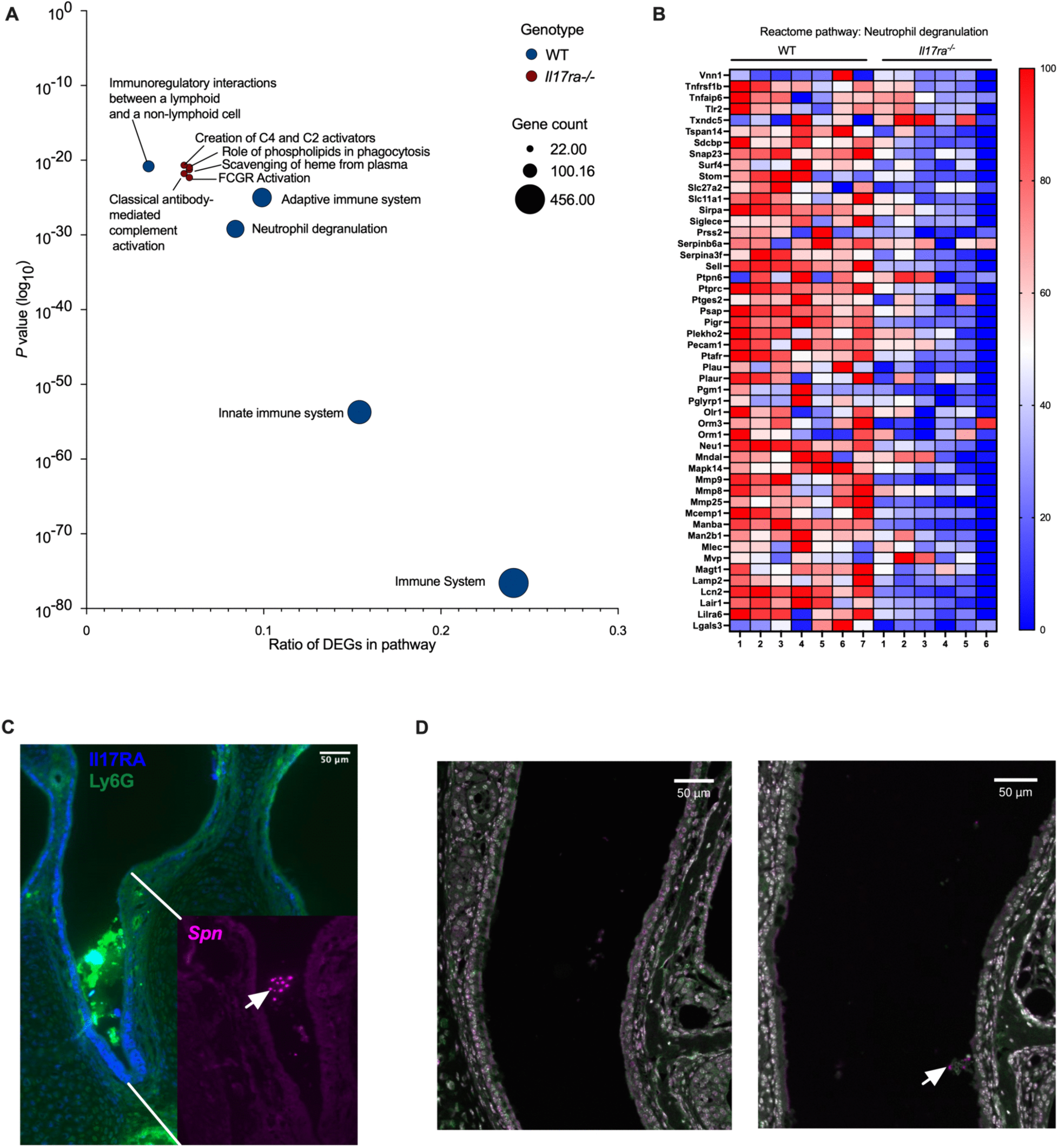
Colonization induces robust immune responses in wild-type but not *Il17ra^-/-^* mice. At 21 dpi, mock- and *Spn*-infected WT and *Il17ra^-/-^* mice were euthanized and nasal lavage samples were collected for RNA isolation and RNA-sequencing. Differentially expressed genes (DEGs) were identified by comparing colonized mice to their respective mock-infected controls within each genotype. The top upregulated genes (p-value <0.05) were analyzed for Reactome pathway enrichment using the Database for Annotation, Visualization, and Integrated Discovery (DAVID) software. **(A)** Top 5 enriched Reactome pathways for Spn-infected WT and *Il17ra*^⁻/⁻^ mice relative to their mock controls. The x-axis represents the gene ratio, and the y-axis shows the log10-transformed *p* value. Circle size corresponds to the number of genes within each pathway. **(B)** Heatmap showing the top 50 most significantly expressed genes within the “Neutrophil Degranulation” pathway. Expression values for *Spn*-infected WT and *Il17ra*^⁻/⁻^ mice were normalized to their respective mock-infected controls and log₂ (x+1) transformed. Individual mice were normalized to the smallest (0%) and largest (100%) value for each gene and presented as a percentage within this range. **(C)** Immunofluorescent images of coronal sections of the mouse nasopharynx collected 7dpi. Sections were stained for IL17RA (blue) and Ly6G (green). The inset is stained for *Spn* 23F capsule (magenta). Scale bar: 50uM **(D)** Immunofluorescent images of coronal sections of the mouse nasopharynx collected 21dpi. Sections were stained for DAPI (white), *Spn* 23F capsule (magenta) and Ly6G (neutrophils; green). Scale bar: 50uM

To gain a better understanding of what neutrophils are doing at the site of colonization, we next analyzed tissue sections of the nasal cavity of colonized WT mice. During early colonization (7 dpi), before the initiation of clearance **(Fig 1A)**, there was a robust influx of Ly6G^+^ neutrophils into the lumen of the nasal cavity at sites where colonizing *Spn* were observed **(Fig 2C)**. In addition, staining for IL-17RA was evident on cells lining the respiratory epithelium as well as the Ly6G^+^ neutrophils accumulating in the lumen, further linking IL-17RA signaling to the local immune response **(Fig 2C)**. Interestingly, at 21 dpi, when clearance is initiated, collections of neutrophils associated with *Spn* in the nasal lumen were no longer seen in WT mice **(Fig 2D)**. The few neutrophils seen in the submucosa were not associated with colonizing *Spn* and did not differ between mock and colonized WT mice **(Fig 2D)**.

### Reduced neutrophil replenishment is associated with IL17RA-deficiency

To assess neutrophil dynamics during *Spn* colonization, neutrophil turnover was quantified at the onset of bacterial clearance. The thymidine analog 5-ethynyl-2′-deoxyuridine (EdU), was administered systemically five days prior to analysis, and EdU incorporation was assessed by flow cytometry in neutrophils isolated from blood and nasal tissue at 21 dpi. *Spn* colonization was associated with increased EdU staining in neutrophils in both nasal tissue and blood **(Fig 3A)**. This observation suggested that neutrophil turnover at the local site of infection was driving proliferation of neutrophil progenitors systemically, with mobilization of these cells to the nasal tissue. However, no difference in the percentage of neutrophils in nasal tissue was observed between mock- and *Spn*-infected WT mice at 21 dpi **(Fig 3B)**. Together with the findings from the EdU experiments, this indicated that the of loss of neutrophils at the site of infection was fully compensated for by the continued recruitment of newly generated neutrophils from the circulation.

**Fig 3:**
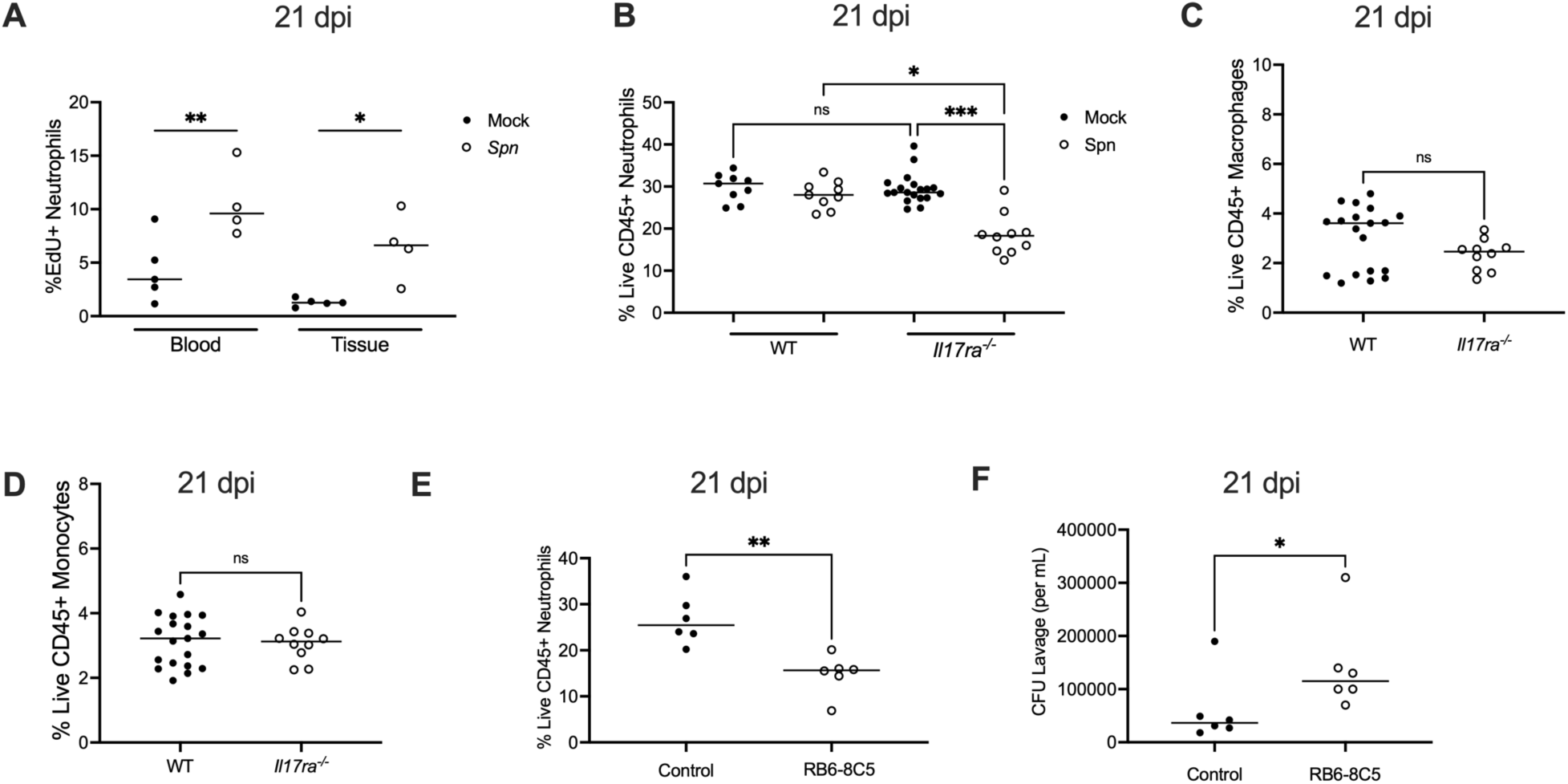
Reduced neutrophil replenishment is associated with IL17RA-deficiency. **(A)** Frequency of EdU⁺ neutrophils (singlet, live CD45⁺ CD11b⁺ Ly6G⁺) in nasal tissue and blood from mock- and *Spn*-infected WT mice injected intraperitoneally with EdU at 16 dpi and euthanized at 21 dpi. Each circle represents an individual mouse and the horizontal line denotes the median for each group. Statistical significance determined using one-way ANOVA with a Sidak’s post-hoc multiple comparisons test. Significance is designated by * = p < 0.05; ** = p < 0.01. **(B)**Frequency of neutrophils (Singlet live CD45⁺ CD11b^+^ Ly6G⁺) in nasal tissue from mock- and *Spn*-infected WT and *Il17ra*^⁻/⁻^ mice at 21 dpi. Each circle represents 2 mice pooled together and the horizontal line denotes the median for each group. Statistical significance determined using one-way ANOVA with a Kruskal-wallis post-hoc multiple comparisons test. Significance is designated by * = p < 0.05; *** = p < 0.001; ns = not significant. **(C)** Frequency of macrophages (Singlet live CD45⁺ F4/80⁺ CD11b⁺) and (**D**) monocytes (Singlet live CD45⁺ Ly6C⁺ CD11b⁺) in nasal tissue from *Spn*-infected WT and *Il17ra*^⁻/⁻^ mice at 21 dpi. Each circle represents 2-3 mice pooled together and the horizontal line denotes the median for each group. Statistical significance determined using unpaired Students *t* test. Significance is designated by ns = not significant. **(E)** Frequency of neutrophils (Singlet live CD45⁺ CD11b^+^ Ly6G⁺) in nasal tissue from WT mice treated with a control mAB or anti-Gr-1 (clone RB6-8C5) mAB antibody at 21 dpi. Each circle represents an individual mouse, and the horizontal line denotes the median for each group. Statistical significance determined using unpaired Students *t* test. Significance is designated by ** = p < 0.01. **(F)** Colonization density of mice treated with control mAb or anti-Gr-1(RB6-8C5) mAb at 21 dpi. Each circle represents an individual mouse, and the horizontal line denotes the median for each group. Statistical significance determined using Mann-Whitney test. Significance is designated by * = p < 0.05.

While mock-infected *Il17ra^-/-^* mice showed similar percentages of neutrophils to WT, colonized *Il17ra^-/-^* mice showed a significantly reduced neutrophil percentage compared to other groups **(Fig 3B)**. This result suggested that during homeostasis neutrophil recruitment itself is not impaired in the absence of IL-17RA signaling. Rather, IL-17RA signaling is required to sustain neutrophil levels which are otherwise diminished without adequate replenishment during colonization. A further implication is that the signature for neutrophil degranulation in the RNA-seq analysis could represent fewer neutrophils rather than diminished function. Unlike neutrophils, we did not observe any significant differences in the macrophage or monocyte populations between WT and *Il17ra^-/-^* mice during colonization **(Fig 3C and 3D)**. This suggests that among the myeloid cell populations, the effects of IL-17RA signaling in response to late colonization are most apparent on neutrophils.

### Neutrophils contribute to clearance during late colonization

To directly test the role of neutrophils in *Spn* clearance, myeloid cells were depleted in WT mice by systemic administration of the anti-Gr-1 mAb (clone RB6-8C5) **(Fig 3E)** [20]. This treatment was chosen over more specific depletion strategies to maximize the effect on mucosal neutrophils. As expected, mAb RB6-8C5 was highly effective in depleting both neutrophils and monocytes in the blood **(S1 Fig)**. In contrast, in nasal tissue, neutrophils were partially depleted (∼40%), while monocyte numbers were unaffected **(Fig 3E and S2 Fig)**. This depletion in tissues could only be maintained for one week, presumably before the generation of anti-RB6-8C5 antibodies limited its effectiveness. Despite these limitations, at 7 days post-depletion, mice colonized for 21 days that received the RB6-8C5 mAb exhibited significantly higher colonization densities compared to isotype control-treated animals **(Fig 3F)**. This result confirmed that neutrophils contribute to *Spn* clearance during late colonization.

### Neutrophil loss is not due to pneumolysin, reduced chemokine-driven recruitment, or direct IL-17RA signaling on neutrophils

We next investigated the mechanism underlying the loss of neutrophils in *Il17ra^-/-^* mice. First, we asked if the loss of neutrophils is attributed to pneumolysin, the pore forming toxin expressed by *Spn* that mediates the lysis of host cells [21], [22]. When *Il17ra^-/-^*mice were colonized with an isogenic pneumolysin-deficient strain of *Spn*, the loss of neutrophils was still evident **(Fig 4A)**. This indicated that the IL-17RA signaling-dependent loss of neutrophils was independent of *Spn*’s sole toxin. Additionally, fewer neutrophils in the absence of Il17RA-signaling was not due to apoptotic cell death as determined by annexin V staining **(Fig 4B)**.

**Fig 4:**
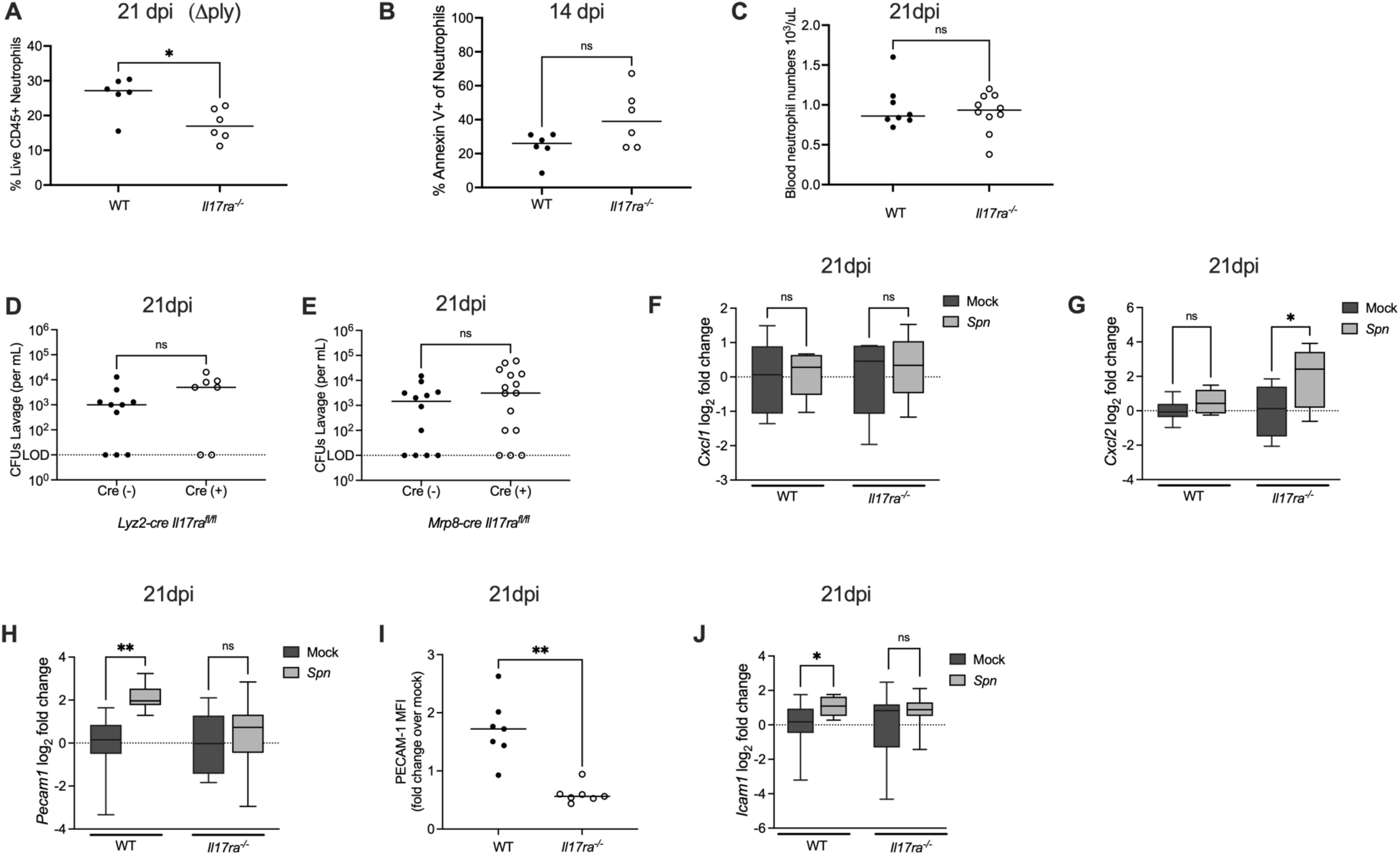
IL-17RA-dependent upregulation of adhesion molecules correlates with increased neutrophil influx during late colonization. **(A)** Infant WT and *Il17ra^-/-^* mice were colonized with a pneumolysin-deficient strain of *S.pneumoniae* (*Δply*) and euthanized at 21 dpi. Frequency of neutrophils (Singlet live CD45⁺ CD11b⁺ Ly6G⁺) among total live CD45⁺ cells in colonized WT and *Il17ra*^⁻/⁻^ mice at 21 dpi. Each circle represents 2 mice pooled together and the horizontal line denotes the median for each group. Statistical significance determined using Mann-Whitney test Significance is designated by * = p < 0.05. **(B)** Frequency of Annexin V+ neutrophils (Singlet live CD45⁺ CD11b⁺ Ly6G⁺) in nasal tissue from *Spn*-infected WT and *Il17ra*^⁻/⁻^ mice at 14 dpi. Each circle represents 2 mice pooled together and the horizontal line denotes the median for each group. Statistical significance determined using unpaired Students *t* test. Significance is designated by ns = not significant. **(C)** Blood neutrophil counts in *Spn*-infected WT and *Il17ra*^⁻/⁻^ mice at 21 dpi. Each circle represents an individual mouse and the horizontal line denotes the median for each group. Statistical significance determined using Mann-Whitney test. Significance is designated by ns = not significant. **(D-E)** Colonization densities of *Lyz2-cre Il17ra^fl/fl^* and *Mrp8-cre Il17ra^fl/fl^* mice colonized as infants with serotype 23F *S.pneumoniae.* Mice were euthanized 6 weeks post-infection to assess colonization levels. Each circle represents an individual mouse and the horizontal line denotes the median for each group. Statistical significance determined using Mann-Whitney test. Significance is designated by ns = not significant. **(F-H)** At 21 dpi, mock- and *Spn*-infected WT and *Il17ra*^⁻/⁻^ mice were euthanized and RNA isolated from nasal lavages was used to quantify *Cxcl1, Cxcl2 and Pecam1* expression by qRT-PCR. Expression was normalized to *Gapdh* and calculated as fold change relative to mock-infected controls. Each group represents 5–6 mice and data are presented as a box-and-whisker plot showing the median, interquartile range, and min–max values. Statistical significance determined using a one-way ANOVA with a Sidak’s post-hoc multiple comparisons test. Significance is designated by * = p < 0.05; ** = p < 0.01; *** = p < 0.001; ns = not significant. **(I)** Median fluorescence intensity (MFI) of PECAM-1 on endothelial cells (CD45^-^ CD31^+^) isolated from nasal tissue of mock- and *Spn*-infected WT and *Il17ra^-/-^*mice at 21 dpi. Data are shown as fold change relative to mock-infected controls. Each circle represents pooled samples from two mice. Statistical significance determined using Mann-Whitney test. Significance is designated by ** = p < 0.01. **(J)** At 21 dpi, WT and *Il17ra*^⁻/⁻^ mice were euthanized and RNA isolated from nasal lavages was used to quantify *Icam1* expression by qRT-PCR. Expression was normalized to *Gapdh* and calculated as fold change relative to mock-infected controls. Each group represents 5–6 mice and data are presented as a box-and-whisker plot showing the median, interquartile range, and min–max values. Statistical significance determined using Mann-Whitney test. Significance is designated by * = p < 0.05; ns = not significant.

We then tested whether IL-17RA signaling affects neutrophil availability in the blood or by the induction of chemokines that recruit neutrophils to tissue. Circulating neutrophil numbers were comparable between WT and *Il17ra*^-/-^ **(Fig 4C)**, suggesting no defect in systemic production or availability. We then measured expression of the major murine neutrophil-recruiting chemokines *Cxcl1* and *Cxcl2* in nasal tissue. The expression of these chemokines, neither of which increased during late colonization, was not IL-17RA-dependent **(Fig 4F and 4G)**. These results showed that neither a decreased supply of circulating neutrophils nor impaired chemokine expression explains the tissue-specific loss of neutrophils during colonization.

We next considered the possibility that IL-17RA signaling acts directly on neutrophils to mediate their effects on colonization. To test this, C57BL/6J mice expressing a *LysM-cre* allele (specific for myeloid cells) or a *Mrp8-cre* allele (specific for neutrophils) were bred to mice homozygous for a floxed *Il17ra* allele (IL17RA^flox/flox^). Conditional IL-17RA deficient mice or their Cre-negative littermate controls were infected with *Spn.* Colonization densities were unchanged between Cre-negative and Cre-positive mice in both models at 6 weeks p.i. **(Fig 4D and 4E)**, indicating that IL-17 family cytokines sensed by IL-17RA do not act directly on neutrophils (or other myeloid cells) to facilitate *Spn* clearance.

### IL-17RA-dependent upregulation of adhesion molecules correlates with increased neutrophil influx during late colonization

We then investigated how IL-17RA signaling affects neutrophil trafficking into nasal tissue. In the RNA-seq analysis, one of the pathways upregulated in WT but not *Il17ra^-/-^* mice during colonization was “cell surface interactions at the vascular wall”, suggesting the ability of neutrophils to migrate into the nasal tissue from the circulation was impaired in the absence of IL-17RA signaling. To explore this possibility, we compared the expression of *Pecam-1*, an important adhesion molecule that aids in transendothelial migration by neutrophils to mediate their migration out of the blood and into tissue [23],[24]. qRT-PCR analysis of nasal lavages showed that *Pecam1* expression was significantly upregulated in WT mice following colonization, whereas its levels remained unchanged in *Il17ra^-/-^* mice **(Fig 4H)**. Flow cytometric analysis confirmed reduced PECAM-1 specifically on endothelial cells (CD45^-^CD31^+^) from infected *Il17ra*^-/-^ mice compared to WT controls **(Figure 4I)**. Consistent with this observation, expression of *Icam1*, another adhesion molecule, was also significantly reduced in nasal lavages of *Il17ra^-/-^* mice during colonization **(Fig 4J)**. Together, these findings suggest that expression of adhesion molecules involved in transendothelial migration of neutrophils is affected by loss of the IL-17RA, potentially explaining the requirement of IL-17RA signaling to replenish tissue neutrophils lost during infection.

### IL-1R signaling prevents persistent colonization independently of neutrophil recruitment

We previously showed that the IL-1R signaling pathway contributes to early clearance of *Spn* colonization [17]. To determine whether this effect was sustained and related to the IL-17RA signaling pathway, late clearance in *Il1r1^-/-^* mice was assessed. Similar to *Il17ra^-/-^* mice, *Il1r1^-/-^* infant mice were persistently colonized (≥ 26 weeks p.i.) **(Fig 5A)**. We then determined whether IL-1-signaling was activated during colonization by measuring expression of *Il1a* and *Il1b*, the two main cytokines sensed by the IL-1R. At 21 dpi, expression of *Il1a* was unchanged **(Fig 5B)**. This was not unexpected since *Il1a* is constitutively expressed as an alarmin and is released upon cell death, such as during infection [17]. At 21 dpi, *Il1b* expression was significantly elevated in *Spn*-infected WT mice compared to mock-infected controls **(Fig 5C)**. Given the importance of IL-17RA signaling for sustaining neutrophil populations, we next examined whether IL-1R signaling similarly regulated neutrophil numbers. Flow cytometric analysis revealed no difference in the percentage of neutrophils between colonized WT and *Il1r1^-/-^* mice at 21 dpi **(Fig 5D)**. Consistently, expression of the neutrophil chemokines *Cxcl1* and *Cxcl2* were not lower in the absence of IL-1R signaling at 21 dpi **(Fig 5E and 5F)**. In contrast to *Il17ra^-/-^* mice, increased expression of adhesion molecules *Pecam1* and *Icam1* during *Spn* colonization was maintained in the absence of IL-1R signaling **(Fig 5G and 5H)**. These findings indicate that persistent colonization in *Il1r1^-/-^* mice was not attributable to altered neutrophil trafficking as is the case for *Il17ra^-/-^* mice.

**Fig 5:**
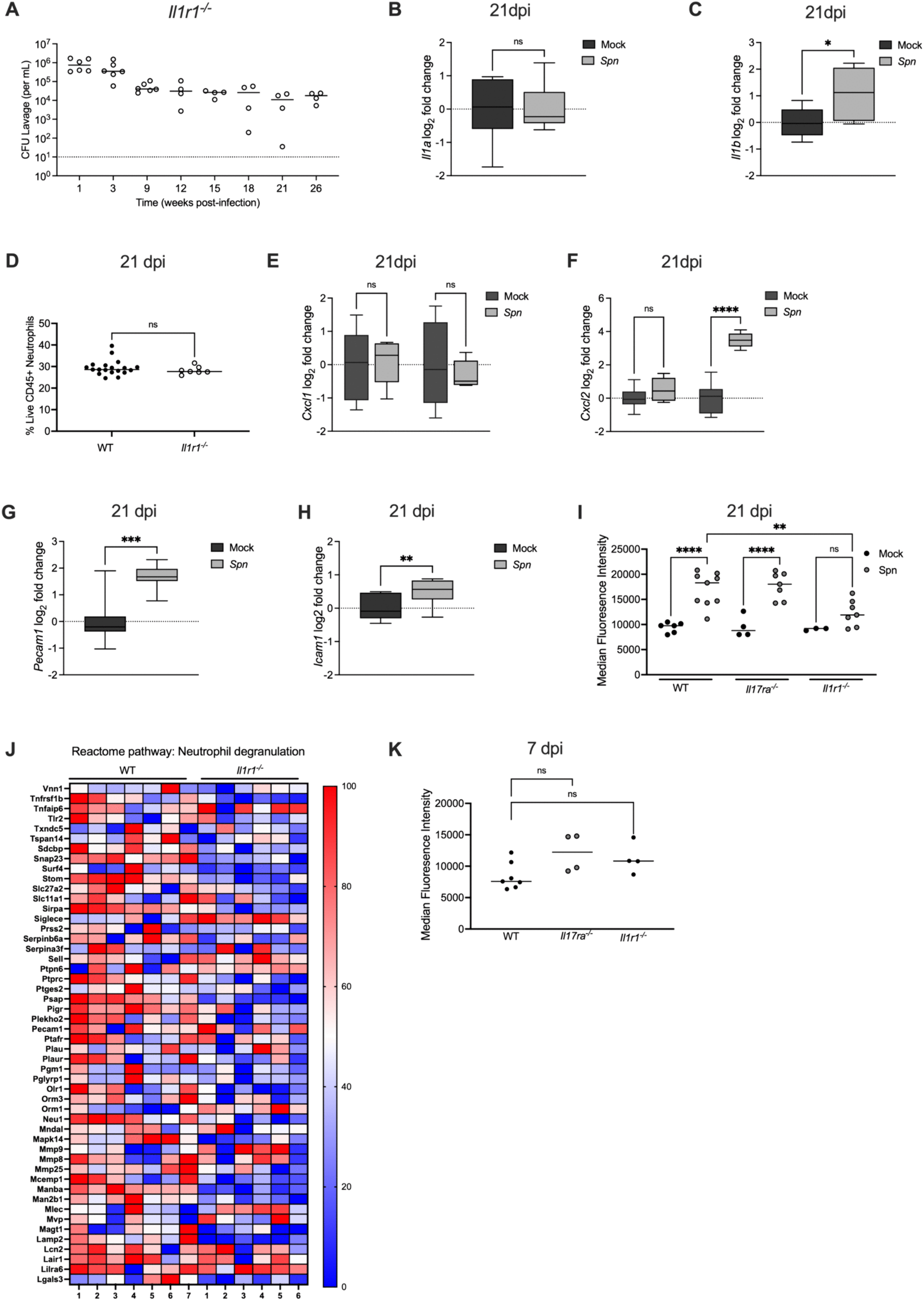
IL-1R signaling prevents persistent colonization independently of neutrophil recruitment. **(A)** *Il1r1^-/-^* mice were colonized intranasally with serotype 23F *S.pneumoniae*. Nasal lavages were collected at the indicated timepoints to quantify colonization density (CFU/mL). Each circle represents an individual mouse and the horizontal line denotes the median for each group. **(B-C)** At 21dpi mock- and *Spn*-infected WT mice were euthanized and RNA isolated from nasal lavages was used to quantify expression of *Il1a* and *Il1b*. Expression was normalized to *Gapdh* and calculated as fold change relative to mock-infected controls. Each group represents 5–6 mice and data are presented as a box-and-whisker plot showing the median, interquartile range, and min–max values. Statistical significance determined using Mann-Whitney test. Significance is designated by * = p < 0.05; ns = not significant. **(D)** Frequency of neutrophils (Singlet live CD45⁺ CD11b⁺ Ly6G⁺) in *Spn*-infected WT and *Il1r1*^⁻/⁻^ mice at 21 dpi. Each circle represents 2 mice pooled together. Statistical significance determined using Mann-Whitney test. Significance is designated by ns = not significant. **(E-F)** At 21 dpi, WT and *Il1r1*^⁻/⁻^ mice were euthanized and RNA isolated from nasal lavages was used to quantify *Cxcl1 and Cxcl2* expression by qRT-PCR. Expression was normalized to *Gapdh* and calculated as fold change relative to mock-infected controls. Each group represents 5–6 mice and data are presented a box-and-whisker plot showing the median, interquartile range, and min–max values. Statistical significance determined using one-way ANOVA with a Sidak’s post-hoc multiple comparisons test. Significance is designated by ** = p < 0.01; ns = not significant. **(G-H)** At 21 dpi, *Il1r1*^⁻/⁻^ mice were euthanized and RNA isolated from nasal lavages was used to quantify *Pecam1* and *Icam1* expression by qRT-PCR. Expression was normalized to *Gapdh* and calculated as fold change relative to mock-infected controls. Each group represents 5–6 mice and data are presented as a box-and-whisker plot showing the median, interquartile range, and min–max values. Statistical significance determined using Mann-Whitney test. Significance is designated by * = p < 0.05; *** = p < 0.001. **(I)** Median fluorescence intensity (MFI) of CD11b expression on neutrophils (Singlet live CD45⁺ CD11b⁺ Ly6G⁺) isolated from nasal tissue of mock- and *Spn*-infected WT, *Il17ra*^⁻/⁻^ and *Il1r1*^⁻/⁻^ mice at 21 dpi. Each circle represents pooled samples from two mice. One-way ANOVA with a Sidak’s post-hoc multiple comparisons test. Significance is designated by ** = p < 0.01; *** = p < 0.0001; ns = not significant. **(J) Heatmap showing the top 50 most significantly expressed genes within the “Neutrophil Degranulation” pathway.** Expression values for *Spn*-infected WT and *Il1r1*^⁻/⁻^ mice were normalized to their respective mock-infected controls and log₂ (x+1) transformed. Individual mice were normalized to the smallest (0%) and largest (100%) value for each gene and presented as a percentage within this range. **(K)** Median fluorescence intensity (MFI) of CD11b expression on neutrophils (Singlet live CD45⁺ CD11b⁺ Ly6G⁺) isolated from nasal tissue of mock- and *Spn*-infected WT, *Il17ra*^⁻/⁻^ and *Il1r1*^⁻/⁻^ mice at 7 dpi. Each circle represents pooled samples from two mice. One-way ANOVA with a Sidak’s post-hoc multiple comparisons test. Significance is designated by ns = not significant.

### IL-1R signaling is required for neutrophil activation during colonization

Seeing that IL-1R signaling did not impact neutrophil percentages, we asked whether it affects neutrophil activation. To test this, we measured CD11b expression on neutrophils by flow cytometry. Importantly, surface expression of CD11b serves as a marker of neutrophil activation, and CD11b forms a heterodimer with CD18 to generate CR3, the iC3b receptor that mediates phagocytosis of complement opsonized *Spn* [25], [26]. At 21 dpi, both WT and *Il17ra^-/-^* mice displayed significantly increased mean fluorescence intensity (MFI) of CD11b on neutrophils, whereas upregulation of CD11b during infection was attenuated in *Il1r1^-/-^* mice **(Fig 5I)**.

Surprisingly, the MFI of CD11b was low (compared to **Fig 5I**) on neutrophils across WT, *Il17ra^-/-^*, and *Il1r1^-/-^* mice at 7 dpi, suggesting that neutrophils during are not activated during early infection, which could explain why the initial presence of neutrophils **(Fig 2C)** is not completely effective at clearance **(Fig 5K)**. The surface expression of CD11b on neutrophils is regulated by inflammatory signals through mobilization from stored granules [27]. To further evaluate whether IL-1R signaling influences neutrophil effector function, we carried out RNA-seq comparing *Spn*-infected WT and *Il1r1^-/-^*mice at 21 dpi and specifically looked at the neutrophil degranulation pathway. At 21 dpi, expression of this pathway was significantly reduced in colonized *Il1r1^-/-^* mice compared to WT controls, consistent with impaired activation of neutrophils **(Figure 5J)**. Together, these results demonstrate that, unlike IL-17RA signaling which maintains neutrophil numbers, IL-1R signaling is associated with the activation and functional programming of neutrophils that is necessary to promote late clearance.

### IL-1R and IL-17RA signaling pathways have nonredundant contributions to clearance

To define the relationship between IL-1R and IL-17RA signaling during colonization, we generated double knockout mice lacking both *Il1r1* and *Il17ra*. We colonized these mice and assessed clearance at 21 dpi. Double knockout mice exhibited significantly higher colonization densities in comparison to either single knockout genotype **(Fig 6A)**. This indicated that the two pathways provide independent and nonredundant contributions to clearance. We next asked whether loss of one pathway influenced activation of the other. By qRT-PCR, we found that during colonization, *Il17a* expression was increased rather than diminished in *Il1r1^-/-^* mice **(Fig 6B)**, and *Il1b* expression was elevated rather than decreased in *Il17ra^-/-^*mice **(Fig 6C)**. This data indicates that disruption of one pathway does not impair activation of the other. All together these observations reveal that both pathways use distinct mechanisms to aid in clearance. In the context of our earlier results, this suggests that the IL-17RA signaling pathway maintains neutrophil numbers during colonization, while the IL-1R signaling pathway activates neutrophils during colonization. Both proper neutrophil trafficking and function are required to prevent persistent *Spn* colonization.

**Fig 6:**
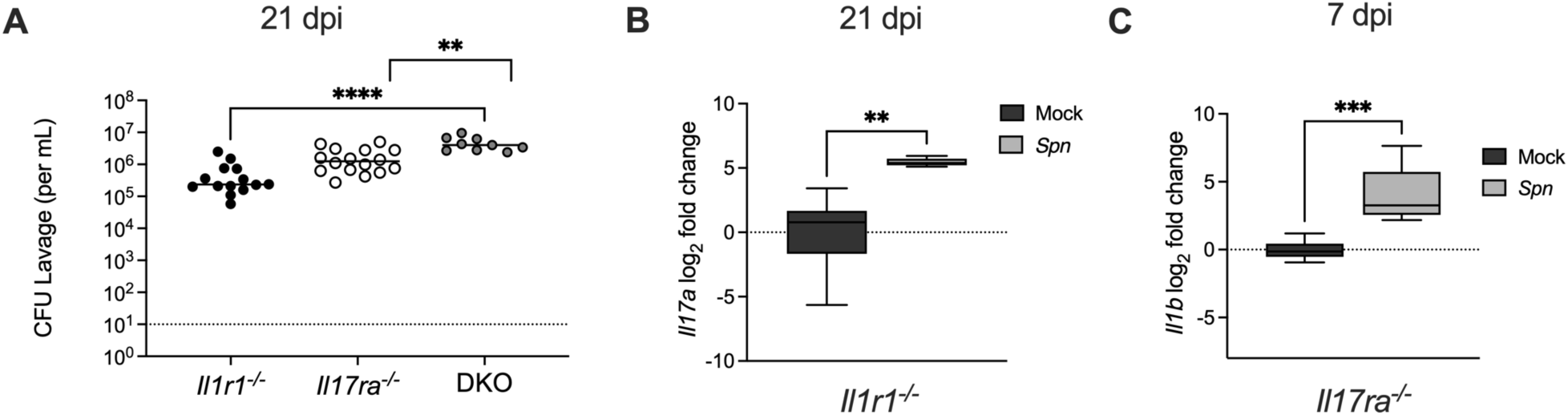
IL-1R and IL-17RA signaling pathways have nonredundant contributions to clearance. **(A)** Double knockout mice (DKO), *Il1r1^-/-^*and *Il17ra^-/-^*, were generated and colonized with serotype 23F *S.pneumoniae*. Colonization density from *Il1r1*⁻/⁻, *Il17ra*⁻/⁻ and DKO mice was assessed at 21 dpi and is shown as CFU/mL. Significance is designated by ** = p < 0.01; **** = p < 0.0001. **(B)** At 21 dpi *Il1r1*^⁻/⁻^ mice were euthanized and RNA isolated from nasal lavages was used to quantify *Il17a* expression by qRT-PCR. Expression was normalized to *Gapdh* and calculated as fold change relative to mock-infected controls. Each group represents 5–6 mice and data are presented a box-and-whisker plot showing the median, interquartile range, and min–max values. Statistical significance determined using Mann-Whitney test. Significance is designated by ** = p < 0.01. **(C)** At 7 dpi *Il17ra*^⁻/⁻^ mice were euthanized and RNA isolated from nasal lavages was used to quantify *Il1b* expression by qRT-PCR. Expression was normalized to *Gapdh* and calculated as fold change relative to mock-infected controls. Each group represents 5–6 mice and data are presented a box-and-whisker plot showing the median, interquartile range, and min–max values. Statistical significance determined using unpaired Students *t* tests. Significance is designated by * = p < 0.05.

## Discussion

Colonization of mucosal surfaces by opportunistic pathogens is generally transient but may be prolonged. While many studies examine how mucosal immunity is induced, few have focused on downstream events that result in eventual clearance and prevent persistent infection [28]. In the case of *Spn*, a leading respiratory pathogen, the expression of a thick layer of capsular polysaccharide greatly delays clearance suggesting that effective host defense requires overcoming its antiphagocytic properties [29],[30]. Our observation of persistent colonization in the absence of IL-17RA-signaling allowed us to look at how the Th17 response and the expression of IL-17-family cytokines mediate mucosal clearance of an encapsulated organism. Our findings provide novel insight into the mechanisms of IL-17RA-signaling that act on neutrophil-mediated defense of mucosal surfaces.

Well before the initiation of the Th17 response there is an influx of neutrophils to the site of infection in the nasal lumen. This early, acute inflammatory response, however, is transient and insufficient for complete clearance. Rather, our findings demonstrate the importance of a sustained presence of neutrophils over many weeks and this requires IL-17RA-signaling. Because our model looks at a prolonged infection, we were able to parse out these later events when IL-17RA-signaling promotes neutrophil-mediated clearance. We excluded a direct effect of IL-17RA-signaling on neutrophils (or myeloid cells) in mediating clearance. When clearance is initiated, coinciding with onset of the Th17 response, there is no increase in neutrophil numbers at the site of infection in nasal tissue. In the absence of IL-17RA-signaling, however, fewer neutrophils are detected in the setting of *Spn* infection. This suggests that neutrophils are being depleted by the infection, and that IL-17RA-signaling is required to compensate for their loss. We also showed that *Spn* colonization drives neutrophil proliferation systemically, indicating that their loss by the mucosal infection is compensated for by Il-17RA signaling. Without IL-17RA- dependent neutrophil replenishment, colonization is persistent. Our data, moreover, supports the importance of IL-17RA-signaling in neutrophil trafficking through the upregulation of adhesion molecules, including on endothelial cells, that could promote neutrophil transmigration from the circulation at sites of infection [31],[32]. Our *in vivo* model is not able to establish whether the increased expression of adhesion molecules is a direct or indirect effect of IL-17RA-signaling. In adult mice with more accelerated clearance, depletion of IL-17A was previously shown to be sufficient to block the CCL2-mediated recruitment of monocytes/macrophages contributing to the engulfment of *Spn* along the nasal lumen [8]. In the current study using infant mice, in which induction of IL17A is delayed (manuscript in preparation), we were unable to detect an IL-17RA-dependent recruitment of monocytes/macrophages in nasal tissue. Additionally, unlike in adult mice, clearance was unaffected in *Ccr2*^-/-^ infant mice with impaired monocyte/macrophage recruitment (**S3 Fig**)[33]. In addition, IL-17RA-signaling, in particular, is known to restructure the murine nasal microbiome [34]. Therefore, in inherent caveat of our approach studying colonization is the diverse microbiome of the upper respiratory tract, which could differ in immunocompromised hosts, potentially affecting *Spn* interactions with other members of the flora.

Since colonization is also more prolonged without IL1R1-signaling, we examined its effects on neutrophils and how these two signaling pathways might converge [17]. Persistent colonization, also seen in the absence of IL1R1-signaling, associated with distinct aspects of neutrophil biology. IL1R1-signaling was not required for neutrophil trafficking or increased expression of adhesion molecules, but correlated with enhanced neutrophil activation as demonstrated by an increased level of CD11b integrin on the neutrophil surface, and increased expression of the neutrophil degranulation pathway. Rapid mobilization of CD11b to the neutrophil surface during activation enhances their phagocytic capabilities [27]. We have previously shown how neutrophil-mediated host defense against invasive *Spn* infection requires CD11b and is highly sensitive to differences in the level of this receptor on the cell surface [26]. Because of the importance of the CD11b integrin in recognition and uptake of complement opsonized targets, our finding suggests that phagocytic clearance by neutrophils may be impaired during colonization in the absence of IL-1R-signaling. We previously described how *Spn* uptake is a catastrophic event for both the bacteria and the professional phagocyte, which undergoes cell death [14]. This process could account for the loss of neutrophils during infection. A further implication is that due to the turnover of these more functionally activated cells they must be replenished, and this process requires IL-17RA-signaling. Neutrophil activation was not observed during early infection, perhaps accounting for the lack of effective clearance during the first days of colonization. The lack of a temporal association with neutrophil activation could be because IL-1R-signaling is necessary but not sufficient to stimulate neutrophil activation. The effect of IL-1R-signaing on neutrophil activation in our report might also be indirect as previously proposed [35], although in another report IL-1β blockade reduced CD11b integrin expression *in vivo* [36]. Alternatively, the cellular release of IL-1a, the main IL-1-family cytokine triggering clearance, might be delayed [17]. Clearance of colonization is also delayed during infection with a mutant lacking the pore-forming *Spn* toxin, pneumolysin [17]. This partially phenocopies infection of the WT strain in the absence of IL-1R-signaling. We were, however, unable to detect an effect of pneumolysin on the IL-1R-mediated increase in neutrophil surface CD11b, suggesting that other bacterial factors also contribute to the activation of neutrophils.

Our *in vivo* study shows that IL-17RA- and IL-1R-signaling pathways have non-redundant independent roles in promoting clearance of *Spn* colonization. Although both pathways are required to prevent persistent colonization, they act through distinct mechanisms with IL-17RA-signaling maintaining the neutrophil population by supporting neutrophil replenishment, while IL-1R-signaling ensures these neutrophils are activated in a way that facilitates *Spn* clearance. The timing of clearance, which determines the duration of carriage, might depend on the two processes converging. Both proper neutrophil trafficking and function are required to prevent persistent colonization.

## Materials and Methods

### Mouse Studies

C57BL/6J, *Il1r1^⁻/⁻^*, LysM-cre and MRP8-Cre-ires/GFP were obtained from the Jackson Laboratories. *Il17ra^-/-^*mice were generously provided by Amgen under a material transfer agreement. Conditional *Il17ra* knockout lines were generated by crossing LysM-Cre and Mrp8-Cre mice with *Il17ra* floxed mice generously provided by Dr. Shruti Naik (Icahn School of Medicine at Mount Sinai). All mouse lines were maintained in an ABSL-2 facility and all studies were approved by the Institutional Animal Care and Use Committee (IACUC) of the New York University Grossman School of Medicine.

Infant mice 4-7 days of age were colonized with 10^3^−10^4^ CFU of *Spn* in 3 μl PBS by atraumatic intranasal instillation without anesthesia. Adult mice 6–10 weeks old were colonized with 10^6^ CFU of *Spn* in 10 μl PBS by intranasal instillation. At the time points indicated following infection, mice were euthanized by CO2 asphyxiation followed by cardiac puncture. The trachea was lavaged retrograde with 400μl of sterile PBS collected from the nares to determine upper respiratory tract (URT) colonization density of *Spn*. For gene expression studies, this lavage was followed up with a second URT lavage with 600μL lysis buffer (RLT lysis buffer (Qiagen) + 1% β-mercaptoethanol) to obtain host RNA from the URT. For histopathology studies, heads and the skin covering the heads were removed following euthanasia. Heads were briefly rinsed in cold dPBS before being transferred into cold 4% paraformaldehyde for 48-hour fixation at 4 °C. Heads were washed in cold PBS for 30 minutes and were transferred into EDTA (0.5M, pH 8) for decalcification. Samples were incubated in EDTA for 7 days (gently shaking, 4°C); EDTA was replaced daily. Heads were washed in cold PBS for 30 minutes, incubated in 50% ethanol for 30 minutes and then were stored in 70% ethanol.

### Bacterial Strain and Infection

A streptomycin-resistant serotype 23F (P2499) pneumococcal strain derived from a clinical isolates was used in this study [37][38]. We also used an isogenic strain with an inframe, unmarked deletion of *ply* (P1726) and its construction has been previously described[39][40]. All strains were grown in tryptic soy (TS) broth in a 37°C water bath to an optical density of 1.0 at 620 nm (OD620). Once at desired OD, bacteria were pelleted and resuspended at the desired density in PBS for colonization studies. For quantitative culture, serial dilutions were plated on TS agar supplemented with 200 μg/mL streptomycin and 100uL of catalase (30,000 U/ml; Worthington Biomedical) and incubated overnight at 37°C with 5% CO₂. CFU were enumerated and expressed as CFU per milliliter of lavage fluid.

### RNA Isolation and Quantitative RT-PCR

For gene expression analysis, tracheal lavages were performed with sterile PBS followed by 600 µL of lysis buffer. RNA was isolated from the lysis buffer lavage using the RNeasy Mini Kit (Qiagen) according to the manufacturer’s instructions. RNA concentration and purity were assessed using a NanoDrop, and samples were stored at −80 °C until further use. cDNA was synthesized using the High-Capacity cDNA Reverse Transcription Kit (Thermo Fisher Scientific). Quantitative RT-PCR was performed using SYBR Green Master Mix (Bio-Rad) on a CFX96 real-time PCR system (Bio-Rad). Gene expression was normalized to *Gapdh* and analyzed using the ΔΔCt method. Primer sequences for *Il17a*, *Pecam1*, *Icam1*, *Cxcl1*, *Cxcl2*, and *Gapdh* are available upon request.

### Flow Cytometry

Following euthanasia by CO2 asphyxiation and cardiac puncture, nasal tissue cells were isolated by removing the lower jaw and tip of the nose and peeling back the hard palate to expose the nasal cavity. The bilateral nasal turbinates were collected along with any remaining tissue in the cavity. Tissues were placed in 1 mL of ice-cold DMEM and digested in 1 mL of DMEM containing collagenase II (200 U/mL, ThermoFisher) and DNase I (15 U/ml; Sigma-Aldrich) for 1 h and 10 min at 37 °C with gentle agitation. Digests were filtered through 70 µm strainers, washed with ice-cold dPBS, and centrifuged at 300 × *g* for 10 min at 4 °C. Pellets were treated with ACK lysing buffer for 5 min at room temperature, washed, and filtered through 40 µm strainers. Cells were stained with LIVE/DEAD Fixable Blue Dead Cell Stain Kit (ThermoFisher). After Live/Dead staining, cells were washed with dPBS and re-suspended in 50 μl of fluorescence-activated cell sorter (FACS) buffer (dPBS containing 1% BSA and 2 mM EDTA), followed by staining with a FC receptor-blocking antibody, anti-CD16/32 (BioLegend; clone 93) for 10 mins at 4 °C. Afterwards, cells were stained with anti-CD45 APC-Cy7 (BD; clone 30-F11), anti-F4/80 PE (BioLegend; clone BM8), anti-CD11b V450 (BD; clone M1/70), anti-Ly6G PerCp/Cy5.5 (BD; clone 1A8), anti-Ly6C AF700 (Biolegend; clone HK1.4) and anti-CD31 (PECAM-1) (ThermoFisher; clone 390) for 30 minutes at 4°C. Stained cells were fixed in 4% paraformaldehyde and resuspended in FACS for flow cytometric analysis. Compensation controls were prepared using OneComp beads (ThermoFisher). Data were acquired on a BD FACSymphony flow cytometer and analyzed using FlowJo.

### Annexin V Staining

Cells were isolated from nasal tissue and stained for surface markers as described above, followed by Annexin V staining using the eBioscience™ Annexin V Apoptosis Detection Kit (Invitrogen). After staining for neutrophils, cells were washed with the kit-supplied binding buffer and incubated in binding buffer containing FITC-conjugated Annexin V for 15 minutes at room temperature. Cells were then washed with binding buffer, fixed in 4% paraformaldehyde, and resuspended in FACS buffer (DPBS with 1% BSA) for flow cytometric analysis. Data were acquired on a BD FACSymphony flow cytometer and analyzed using FlowJo.

### Neutrophil Turnover

Mock and *Spn*-infected infant mice received an intraperitoneal injection of the thymidine analog 5-ethynyl-2′-deoxyuridine (EdU) (50 mg/kg) at 16 dpi and were euthanized at 21 dpi [41]. Nasal tissue was collected and processed as described above. Blood was collected by cardiac puncture, diluted in 10 mL dPBS, and centrifuged at 1500 rpm for 5 minutes at 4 °C. Cell pellets were treated with ACK lysis buffer for 5 minutes at room temperature, washed, and stained using the LIVE/DEAD Fixable Blue Dead Cell Stain Kit (Thermo Fisher Scientific). Cells were then processed for analysis as described above. EdU incorporation, a measure of proliferation, was assessed using the Click-iT™ EdU Flow Cytometry Assay Kit (Invitrogen, C10420) according to the manufacturer’s instructions. Data were acquired on a BD FACSymphony flow cytometer and analyzed using FlowJo software.

### Tissue Preparation for Immunofluorescence and Microscopy

For the 7 dpi images, tissues were processed as previously described [42],[43],[44]. Briefly, sections were washed with PBS and blocked with anti-CD16/32 (BioLegend; clone 93) in PBS containing 2% donkey serum and 2% FBS for 1 h at room temperature. Primary antibodies included rabbit anti-mouse IL-17RA (Invitrogen), Ly6G-AF488, Ly6G-BV421, and rabbit *Spn* 23F antiserum (Statens Serum Institute). Antibodies were diluted in PBS with 2% donkey serum, 2% FBS, and 0.05% Fc block and incubated for 1 h at room temperature. After washing, slides were stained with donkey anti-rabbit 647 (Invitrogen) for 1 h at room temperature in the dark. Sections were mounted with Immun-Mount (Fisher Scientific) and imaged on a BZ-X800 All-in-One fluorescence microscope (Keyence) using BV LP, eGFP, Cy3/TRITC, and Cy5 filter cubes and a 20× Plan Fluor objective. Whole-section images were stitched using BZ-X800 Analyzer software (Keyence).

For the 21 dpi images, tissues were fixed in 4% PFA at 4°C for 48 hours, dehydrated through graded ethanols and xylene, then infiltrated with paraffin (Paraplast X-tra; Leica; cat. 39603002) on a Leica Peloris II tissue processor and embedded on a Leica Arcadia embedder. Five-µm tissue sections were immunostained on a Leica BondRx autostainer according to the manufacturer’s instructions. Sections were deparaffinized online, treated with 3% H₂O₂ to quench endogenous peroxidase activity, and subjected to antigen retrieval using ER1 (Leica, AR9961; pH 6) or ER2 (Leica, AR9640; pH 9) buffer at 100 °C for 20 or 60 minutes. Depending on the species of the primary antibody, tissues were blocked with Rodent Block M (Biocare, RBM961) followed by Primary Antibody Diluent (Leica, AR93520), or with Primary Antibody Diluent alone, prior to primary antibody incubation. Primary antibodies included Ly6G (BD Biosciences; clone 1A8) and rabbit *Spn* 23F antiserum (Statens Serum Institute). Sections were sequentially incubated with primary antibodies and HRP polymer secondary reagents, followed by HRP-mediated tyramide signal amplification using Opal® fluorophores. Opal® fluorophores included Opal 620 (Akoya Biosciences; FP1495001KT) and Opal 570 (Akoya Biosciences; FP1488001KT). After each staining cycle, bound antibodies were removed by heat-induced retrieval, and the process was repeated for a total of six markers. Sections were counterstained with spectral DAPI (Akoya Biosciences; FP1490) and mounted with ProLong Gold Antifade (Thermo Fisher Scientific; P36935). Multispectral image acquisition was performed on an Akoya Vectra Polaris (PhenoImager HT) system. Slides were scanned at 20x magnification using PhenoImager HT 2.0 software in conjunction with Phenochart 2.0 and InForm 3.0 to generate unmixed whole-slide qptiff images.

### RNA-Sequencing and Pathway Analysis

RNA-sequencing was performed on RNA isolated from nasal lysis buffer lavage samples of mock-infected and colonized mice. RNA was isolated as described above and six to seven replicates per group were subjected to RNA-sequencing. Automated stranded RNA-seq libraries were prepared using poly(A) selection and sequenced on an Illumina NovaSeq 6000 platform. Differential expression analysis was conducted using DESeq2 and genes with an adjusted *p*-value < 0.05 were considered significantly differentially expressed. Differentially expressed genes were analyzed to identify enriched reactome pathways using the online annotation tool DAVID Bioinformatics Resources [45]. Heatmaps were generated from the differentially expressed genes using GraphPad Prism version 10.0.

### Neutrophil Depletion

Mice were treated intraperitoneally with 200μg of rat anti-mouse Gr-1 monoclonal antibody (BioXcell; RB6-8C5; cat. #BE0075) or an isotype control (BioXcell; IgG2b; cat. #BE0090) diluted in 40ul of PBS. The antibodies were administered every three days over a 7-day period for a total of 3 doses. Depletion efficacy was confirmed by flow cytometric analysis of nasal tissue. To assess blood neutrophils and monocytes, blood was harvested through cardiac puncture and collected in an EDTA-coated microtainer tube (BD Biosciences). Samples were analyzed using the Element HT5 Veterinary Hematology Analyzer (Heska) according to the manufacturer’s instructions.

### Statistical Analysis

Data were analyzed using GraphPad Prism version 10.0. Statistical significance was determined using unpaired two-tailed Student’s *t*-test for two-group comparisons or one-way ANOVA for multiple comparisons, as appropriate. All data are presented as individual values with the median indicated, and *p* < 0.05 was considered statistically significant.

## Acknowledgements

We would like to thank the NYU School of Medicine’s Genome Technology Center for library preparation and sequencing. We also thank the Applied Bioinformatics Laboratories at the NYU School of Medicine for providing bioinformatics support and analysis of the data. In addition, we would like to thank Dr. Cynthia Loomis and the experimental pathology core for their assistance with the nasal tissue section preparation and staining. We thank Dr. Chris Hergott for his valuable guidance and assistance with the EdU experiments. Finally, we would like to thank Amgen for their generous donation of the *Il17ra^-/-^* mice. This study was supported by NIH grants to J.N.W. (R01 A150893 and R37 AI38446).

